# Individual Decision-Making Underlying the Tragedy of the Commons

**DOI:** 10.1101/2022.11.29.518377

**Authors:** Megha Chawla, Matthew Piva, Shamma Ahmed, Ruonan Jia, Ifat Levy, Steve W. C. Chang

**Author notes:** Co-first authors (Megha Chawla, Matthew Piva).

## Abstract

Group decision-making is common in everyday life, whether a family is sharing a meal or a corporation is dividing profits. Research in economics on group decision-making has coalesced into a theory known as the tragedy of the commons, which states that resources are inevitably overused when shared by a group. However, even while multiple approaches to mitigating overuse of common resources have been put forward, notable counterexamples to the tragedy of the commons exist such that groups are ultimately able to avoid resource overuse. Development of a computerized paradigm amenable to behavioral modeling and simulation analyses could allow for exploration of whether resources will be overused in a given group of individuals and allow for the rapid testing of behavioral interventions designed to reduce instances of resource overuse. Using a newly developed group decision-making task, we studied how participants made decisions to utilize shared resources for the potential to receive a larger amount of money or conserve resources for a smaller amount of money. Using behavioral modeling, we found that valuation of resource overuse is most impacted only when an exceptionally small amount of resources are remaining. Using computational analyses, we were able to differentiate individual participants by both group earnings and self-reported social attitudes in ways that correlated with their willingness to utilize resources. These results signify the importance of individual differences in group composition regarding the tragedy of the commons, emphasizing the impact of the attitudes and behaviors of individual group members in predicting shared resource use.

## INTRODUCTION

Decision-making among individuals in a group concerning scarce, commonly held resources has long been a prominent question in economics and a common situation in daily life with strong public policy relevance. Group decision-making is involved in a range of tasks, from sharing a pizza to making the commitment to install solar panels on one’s home. Previous academic research on this problem has coalesced into a central theory known as the tragedy of the commons (Boyd et al., 2018; Crowe, 1969; Dawes, 1980; Hardin, 1968). In this scenario, commonly held resources are overused by the group such that the resource pool cannot be replenished and is therefore no longer useful. This idea has intuitive and immediate importance to overpopulation (Hardin, 1968). However, the implications of the tragedy of the commons have reached far more broadly and have been heavily studied. These include applications in international relations (Nordhaus, 2015), environmental conservation (Ansari, Wijen, & Gray, 2013), evolutionary biology (Rankin, Bargum, & Kokko, 2007), water management (Meinzen-Dick, 2007), and even antimicrobial resistance (Levin, 2001). With these frequent applications have come potential solutions to the problem. Authors of an influential paper reported the need for reputation maintenance as a means of dissuading group members from overusing resources (Milinski, Semmann, & Krambeck, 2002). A more recent paper has reported similar findings in chimpanzees, stating that these animals use dominance as a means of overcoming the problems associated with common resource allocation (Koomen & Herrmann, 2018). Other traditional solutions to the tragedy of the commons include governance (Dietz, Ostrom, & Stern, 2003; Ostrom, 1999; Van Vugt, 2009) and private ownership (Hardin, 1968), and a recent report also indicated the importance of reciprocity (Gachter, Kolle, & Quercia, 2017). Still, not all scholars have agreed with the initial implication of the tragedy of the commons, namely that all commonly held resources would invariably be overused by a group of a sufficient size. Many scholars have noted prominent counterexamples, including multiple case studies in which communities have successfully managed vital resources through careful human cooperation (Berkes, Feeny, McCay, & Acheson, 1989; Boyd, et al., 2018; Feeny, Berkes, McCay, & Acheson, 1990).

While the tragedy of the commons has been heavily debated in the fields of economics, law, and policy research, less attention has been paid to this theory in psychology and neuroscience. Perhaps because of this, current approaches used to probe the tragedy of the commons typically examine decision-making at the group level, rather than at the level of individuals that make up the group. Studying decision-making behaviors using a novel, computerized task that recapitulates the tragedy of the commons can be beneficial to this line of enquiry in multiple aspects. First, such studies enable collection of data from particularly large sample sizes drawing from individuals with diverse backgrounds, which can be used to probe how generalizable the implications of the tragedy of the commons are across many varying groups. This in turn allows the study of individual differences in group decision-making, based on differing underlying social attitudes, political affiliations, income, gender, or the presence of neuropsychiatric disorders, for example. Second, this approach presents new opportunities to allow the rapid quantitative assessment of interventions that may curtail resource overuse in certain circumstances. One could test the previously developed methods of overcoming the tragedy of the commons mentioned above as well as explore additional methods that may be helpful. Finally, tasks used in such studies could also be made to be compatible with various neuroimaging approaches, allowing simultaneous task participation and measurement of neural signals to determine the neural underpinnings of scarce resource utilization within a larger group. Previous studies using a similar approach integrating economics, psychology, and neuroscience have yielded important insights related to subjective value in monetary and social decision-making (Clithero & Rangel, 2014; Kable & Glimcher, 2007; Lockwood et al., 2017; Smith, Clithero, Boltuck, & Huettel, 2014; Wang, Smith, & Delgado, 2016).

To jumpstart this line of research, we have developed a novel group decision-making paradigm amenable to behavioral modeling and simulation in order to recapitulate the tragedy of the commons dilemma in an online task that participants can perform outside the laboratory on their personal computing devices. Capitalizing on this new approach, we investigated how participants make decisions in groups between utilizing shared resources to potentially receive a larger amount of money and conserving shared resources to potentially receive a smaller amount of money. As expected, we observed that both the amount of money one could earn for utilizing resources and the amount of resources available affected the decision of whether or not to use resources. In addition, modeling revealed a parabolic decision-making process, such that the available resources impacted participants’ valuation of options most strongly when these resources were exceptionally scarce. Finally, through the obtained data and further simulations based on them, we were able to differentiate individual participants by both group earnings and self-reported social attitudes in ways that corresponded with their willingness to utilize resources. Our findings emphasize the importance of the behaviors and attitudes of individual group members in influencing group outcomes and lay the groundwork for an exciting area of research that aims to develop the psychological basis for the decisions that individuals make within a group grappling with the allocation of scarce resources.

## METHOD

### Data collection

#### Participants

Eighty-three participants provided online consent to take part in a study that they could complete at any time within a 72-hour time window on a personal computing device of their choosing. All 83 participants successfully completed a quiz that tested understanding of task rules just prior to task completion. Three participants were excluded from all analyses for failing to make logical choices on catch trials during the task, making 80 participants the total sample size of our study (27 male, mean age 26.5, s.d. = 7.6). All participants were between 18 and 55 years of age. The study was approved by the Yale School of Medicine Human Investigation Committee, and all participants were paid for their participation. Sample size was chosen to well exceed recent studies using similar behavioral paradigms (Lockwood, et al., 2017). We also designed our task to be amenable to simulation to allow for greater power in certain statistical analyses.

#### Experimental task and questionnaires

All participants were assigned to one of eight groups totaling ten participants in each group. For groups in which a participant had to be excluded for failing to make logical choices during a catch trial, an additional participant was invited to take part in the study and retroactively assigned to a given group. All members of each group were assigned a 72-hour time period in which they could sign in to a virtual environment supported by the Qualtrics web application to complete the group decision-making task as well as specific self-report questionnaires. Once the 72-hour time period had elapsed, one randomly selected trial was chosen to be played out so that participants received additional funds, in addition to the $10 that every participant was given, after successfully completing the study. The choices of all individual group members for that randomly selected trial were tallied, and participants were provided with additional funds depending on their choices. Funds were transferred to participants electronically using PayPal.

The group decision-making task required participants to decide whether or not to utilize common resources in order to possibly receive more money in a given trial. In every trial, participants decided between two options, the baseline offer and alternative offer. The baseline offer stayed constant at $5 and did not involve utilizing a resource. The alternative offer pseudo-randomly varied between $6, $8, $12, $16, and $25, and always involved utilizing one resource. The amount of resources also pseudo-randomly varied across trials and ranged between one and nine available resources for the group to use. All combinations of alternative offer level and resource amount were used to create 45 unique trials that were completed twice by each participant in a randomized order. If resources in a given trial were utilized past zero, no one in the group would receive any additional funds regardless of what offer they chose. If resources in a given trial were not used up past zero, each individual participant received the offer that they chose. For example, if four resources were available and five or more participants chose the alternative offer, no one in the group would receive any additional funds. However, if four or less people were to choose the alternative offer, those that chose the alternative offer would receive that amount while people who chose the baseline option would receive the baseline $5. Importantly, participants were never made aware of the choices that other members of the group made for any trial aside from having the knowledge that a total of 10 individuals were making decisions with the same amount of resources.

In order to ensure that participants were actively making logical choices throughout the task, two catch trials were randomly interspersed in the 90 task trials. Logical choices on both catch trials were required for inclusion in data analyses. One catch trial involved an alternative offer of $3, and thus the baseline offer of $5 was the logical choice for this trial. The other catch trial involved an alternative offer of $12 and ten available resources. As there was no possibility of the group overusing ten resources, the logical choice for this trial was to choose the alternative offer.

Following completion of all task trials, participants completed three questionnaires to assess social attitudes. These behavioral questionnaires included the Self-Report Altruism Scale (Rushton, Chrisjon, & Fekken, 1981), the Empathy Quotient (Baron-Cohen & Wheelwright, 2004), and the Levenson Self-Report Psychopathy Scale (Levenson, Kiehl, & Fitzpatrick, 1995).

### Data analysis

Unless otherwise noted, all tests were two-tailed with the sample size defined by the number of participants included in each analysis. Significance was defined by *P* < 0.05.

#### Choice-averaged analyses

All participants were included in choice-averaged analyses (*N* = 80). To describe the overall choice behavior of the entire group of participants, we averaged the proportion that individuals chose the alternative offer for each distinct trial type. We then calculated the proportion that individuals chose the alternative offer averaged across alternative reward levels and resource amounts. These data were then put through a two-way ANOVA with alternative reward level and resource amount as factors.

#### Modeling analyses

All participants who chose the alternative offer in at least 5% of trials were included in all modeling analyses (*N* = 75). In order to develop a model to describe behavioral decision-making in our task, we employed four different models, including linear (Eq. 1), exponential (Eq. 2), hyperbolic (Eq. 3), and parabolic (Eq. 4) discounting models. The models were conceptualized as follows:

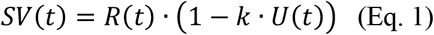

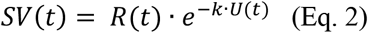

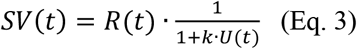

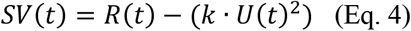

These equations conceptualize subjective value (*SV*) on a certain trial (*t*) as a number that quantifies how likely a participant would be to choose the alternative offer, with higher value indicating a higher probability. This number is modulated by both the alternative offer reward level (*R*) and the amount of resources that are unavailable (*U*), such that *SV* increases as *R* increases but decreases as *U* increases. The models critically diverge in describing how *SV* decreases with higher values of *U*. The parameter *k* is fit individually for each participant and describes the sensitivity of participants to the amount of resources that are unavailable on a given trial, with higher values for *k* indicating a greater sensitivity to the amount of resources available. The softmax function (Eq. 5) was then used to calculate choice probability:

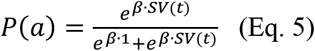

In this equation, *P(a)* refers to the probability that participants would choose the alternative offer. The value of the baseline offer was arbitrarily set to a value of 1. A noise parameter (*β*) defined the stochasticity of each participant’s choices.

Together, a total of four models were tested, the linear, exponential, hyperbolic, and parabolic models. We used Bayesian information criterion (BIC) to compare model performance. We evaluated fit as the sum of BIC values of all individual participants. Additionally, we determined the number of participants in the sample for which each model fit best as determined via BIC. Finally, we examined the example case of $12-alternative offer trials with varying amounts of resources available for best-fitting model of the median participant, the 25^th^-percential participant, and the 75^th^-percentile participant as determined by their individually fitted *k* parameters.

#### Simulation analyses

As the participants in our study were not given any information regarding the other members of their group or any feedback as to the choices that other individuals had made, this uniquely allowed the random matching of individuals to various hypothetically formed groups in order to evaluate task results with an increased sample size. We first used this approach to determine the extent to which participants in our task in these randomly formed groups overused resources past 0 and would therefore receive no reward for these trials. To do this, we created 5000 randomly selected groups drawing from all 80 participants in our study. We then evaluated choice behavior on each trial and determined whether or not resources were overused on each distinct trial type. We then determined the proportion of times that resources were overused for trials with a distinct alternative offer reward level and amount of resources available.

Next, we turned to individual differences in task performance. To evaluate task performance in all 80 participants in our study, we created 1000 groups for each participant that included the participant of interest and nine other randomly selected participants from the sample. We then determined each individual’s task performance, defined as the averaged sum of money earned across all trials, as well as group performance, defined as the averaged sum of money earned across all trials for all other participants in the group excluding the individual of interest. We then determined the correlation between individual and group earnings for all participants using Spearman’s rank correlation. Based on this analysis, we separated individuals as either “low earners,” defined as the bottom 30 individual earners, or “high earners,” defined as the top 30 individual earners. We then compared the individual and group earnings of the low and high earners with a two-way ANOVA, with low versus high earners and individual versus group earnings as factors. We also compared both the proportion of alternative choices and value of the parabolic *k* parameter using Wilcoxon signed-rank tests.

Finally, we created groups that included a certain amount of low or high earners, ranging from groups with no high earners to groups formed exclusively of high earners. To do this, we again randomly sampled participants to form 1000 groups for each distinct group composition. We then evaluated group earnings by calculating the averaged amount of money that each participant in the group would receive if their choices were to be played out. Finally, we also calculated the averaged proportion that each group type overused resources.

#### Individual differences analyses

To determine whether self-reported social attitudes corresponded to behavior in our decision-making task, we performed a principal component analysis on the altruism, empathy, and psychopathy scores of all participants. We then evaluated the correlation between the first principal component score and the individually fitted parabolic *k* parameter of each participant using Spearman’s rank correlation. Three participants were excluded from this analysis for displaying *β* values less than 0.01, and two further participants were excluded for having *k* parameters two standard deviations above the mean. Thus, total sample size for this analysis was *N* = 70. Significance was not affected by the exclusion of the two participants with high *k* parameters, as we observed a significant correlation even with these outlier participants included (*R* = -0.280, *P* = 0.019, Spearman’s rank correlation). Finally, the first principal component scores for the low earner and high earner groups were compared using a Wilcoxon signed-rank test.

#### Code and data availability

The data and codes used in this paper are available and downloadable through GitHub (link to be provided upon publication).

## RESULTS

In the group decision-making task, participants made decisions in each trial between a “baseline offer” of $5 that did not require using any resources and an “alternative offer” of a higher amount of money that always required using one resource (Fig. 1A). The amount of resources in each trial ranged from one to nine resources, while the alternative offer amount was $6, $8, $12, $16, or $25. If each group of 10 participants utilized resources past zero, that is, if more participants in a group chose the alternative offer than there were available resources in a given trial, no one in the group would earn any money, regardless of whether they chose the baseline or alternative offer. However, if resources were not used up past zero, everyone in the group would receive the offer they chose.

**Fig. 1.**
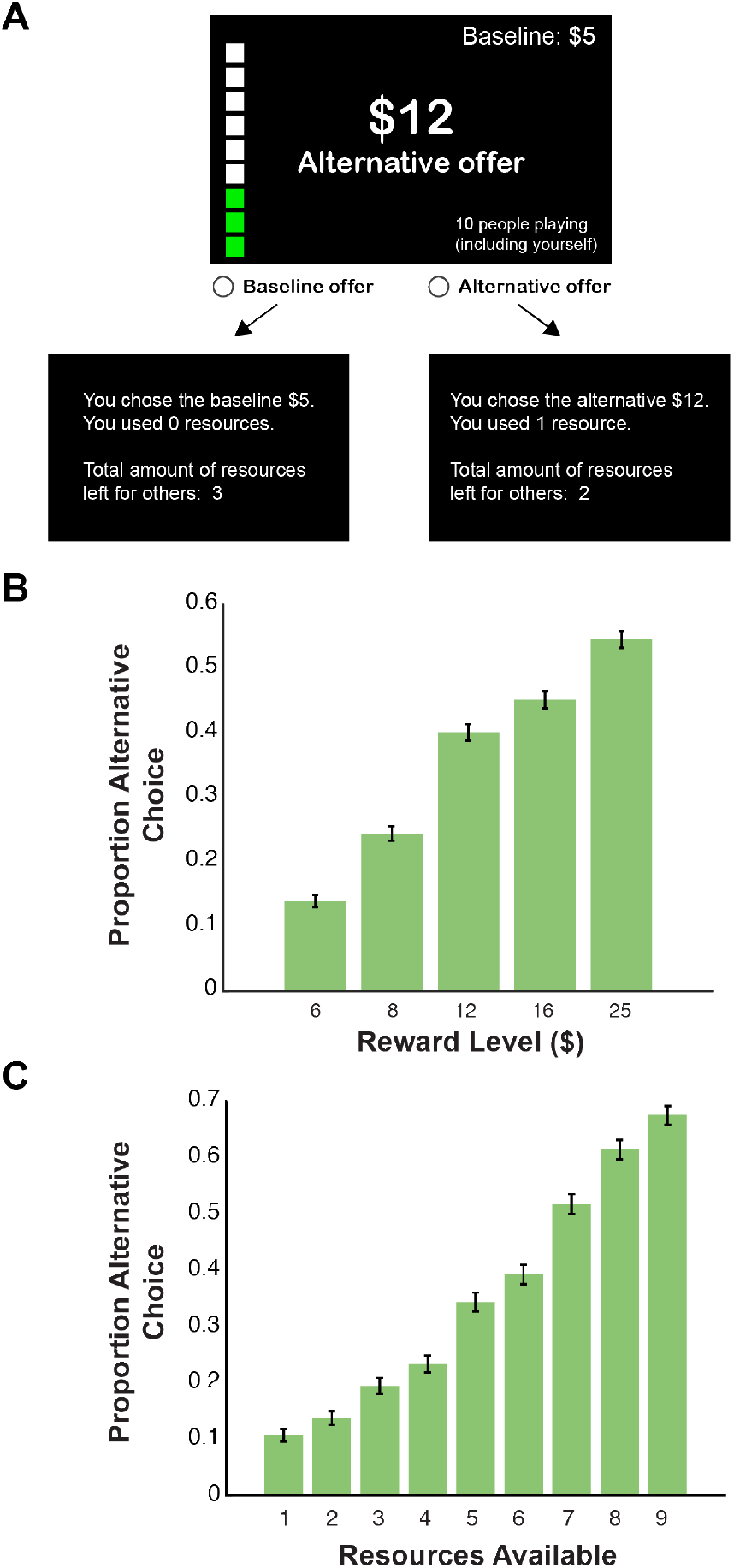
Alternative offer level and resources available influence choice in the group decision-making paradigm. *(A)* An example experimental task progression displaying a $12 alternative offer trial with three available resources for the commons for a 10-participant group. Participants viewed either the alternative or baseline offer feedback screens depending on their choice. *(B)* Proportion of alternative offer choices averaged over all participants for each alternative offer reward level. *(C)* Proportion of alternative offer choices averaged over all participants for each amount of resources available. Error bars indicate SEM. *N* = 80 participants.

First, we determined whether both alternative offer level and resource availability affected choice behavior in our participants to determine whether our task captured the essence of the scarce resource problem. We first calculated the averaged proportion of alternative offer choices for all distinct trial types in the full sample of 80 participants (Fig. 2). Next, we averaged groups of trials by alternative offer level and resource availability (Fig. 1B,C). A two-way ANOVA with alternative offer level and resource availability as factors indicated main effects of both reward level (*F*_4,316_ = 159.0, *P* < 0.001) and resources available (*F*_8,632_ = 143.2, *P* < 0.001). An interaction between these factors was also observed (*F*_8,632_ = 5.3, *P* < 0.001), indicating that more subtle nuances in participant behavior may be further explained by additional analyses using modeling, simulation, and exploration of individual differences.

**Fig. 2.**
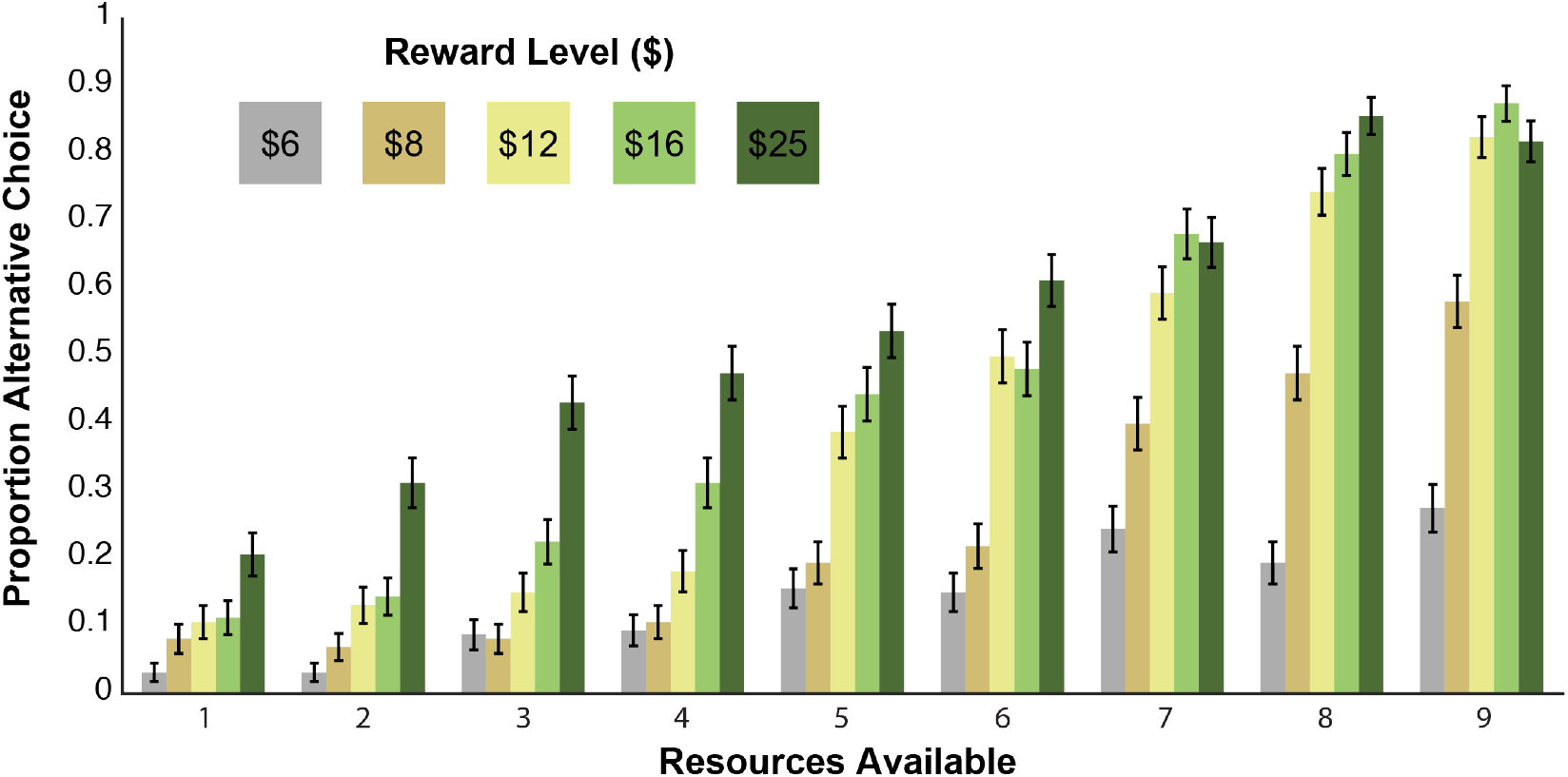
Participant behavior in the group decision-making task is impacted by both alternative offer level and amount of resources available. Averaged proportion of alternative offer choices is displayed for each individual trial type. Error bars indicate SEM. *N* = 80 participants.

### Decision-making under scarce resources follows a parabolic function

Next, we took a modeling approach to describe group decision-making in our task by conceptualizing decisions as being guided by subjective value. In this case, subjective value is a number calculated for each trial, with higher numbers indicating a higher likelihood to choose the alternative offer.

The two quantities that could affect subjective value in our task were the alternative offer level and the amount of resources available. We developed four different models that divergently described lower subjective value in trials that had fewer resources available. Specifically, all the models included the amount of resources that happened to be unavailable in a given trial as a term. For example, in a trial with six resources available, the value of resources unavailable in our model would be four, while in a trial with two resources available, the value of resources unavailable in our model would be eight. Furthermore, subjective value in each model was impacted differently by the amount of resources unavailable. In the linear model (Eq. 1), the subjective value was lowered linearly with the amount of resources that happened to be unavailable in a given trial. In the remaining three models, subjective value was lowered exponentially (Eq. 2), hyperbolically (Eq. 3), or parabolically (Eq. 4), respectively. All of these models contained a free parameter, *k*, which described the sensitivity of each participant to the amount of resources unavailable, with higher values indicating a greater sensitivity to resource availability. Notably, one of these four models may be more consistent with the overutilization of resources noted in the tragedy of the commons scenario (Hardin, 1968). In particular, for the parabolic model, subjective value was substantially affected by resource unavailability only when the amount of resources available was exceptionally low. If members of a given group made decisions using this model, they may have been more likely to overuse middling amounts of resources.

We tested these four models on a sample of participants that chose the alternative offer in at least 5% of trials (*N* = 75) using Bayesian information criterion (BIC). When the individual BIC values for all individual participants were summed, the parabolic model displayed the lowest values and therefore emerged as the best model (Fig. 3A; linear BIC = 3000, exponential BIC = 3241, hyperbolic BIC = 3373, parabolic BIC = 2572). When we evaluated which model fit best at the level of individual participants, the parabolic model described behavior best in the majority of participants (Fig. 3A; linear = 13.3%, exponential = 8.0%, hyperbolic = 25.3%, parabolic = 53.3%).

**Fig. 3.**
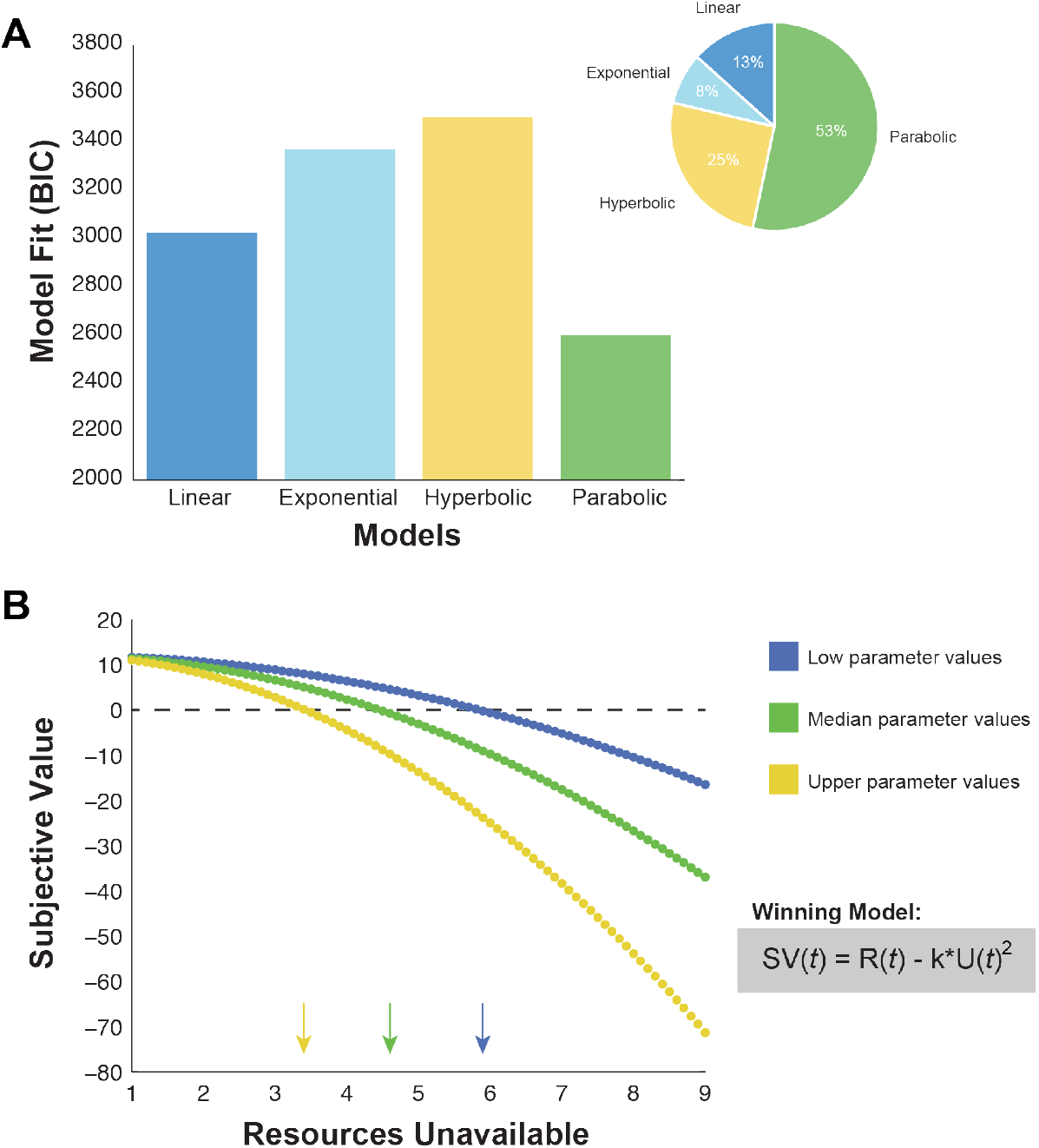
Participant behavior in the group decision-making task follows a parabolic decision-making process. *(A)* Summed Bayesian information criterion (BIC) values across all participants selecting the alternative offer in a minimum of 5% of trials for a total of *N* = 75 participants. Lower BIC values indicate better fit, demonstrating the parabolic model to perform best. The inset shows the percentage of participants who were best fit by each individual model according to BIC values, indicating that the parabolic model performed best for the majority of participants. *(B)* Representative behaviors from the 25^th^ percentile (blue), median (green), and 75^th^ percentile (yellow) participants according to individually fitted *k* parameters for $12 alternative offer trials. Higher levels of subjective value indicate a higher willingness to select the alternative offer. Arrows indicate the indifference point, or point of subjective equality, for each participant, at which one is equally likely to select the alternative and baseline offers. Inset shows the winning parabolic model, where *SV* is subjective value, *R* is the alternative offer level, *U* is the amount of resources unavailable, *t* is a particular trial, and *k* is the individually fitted parameter that describes sensitivity to resource amount, with higher values indicating higher sensitivity.

To determine whether the parabolic model provided realistic predictions of behavior in our task, we selected the median as well as the 25^th^ and 75^th^ percentile participants, as determined by the value of their individually fitted *k* parameters. For trials in which the alternative offer was $12, we plotted the subjective value for trials with varying amounts of resources available (Fig. 3B). For the three representative participants shown in Fig. 3B, the indifference point at which participants were equally likely to choose the alternative or baseline offers was between three and seven resources unavailable, demonstrating that this model made divergent predictions regarding a range of participant behaviors. Together, these analyses indicate that the majority of participants obeyed a parabolic model to make decisions in our task, a pattern of decision-making that could lead to the sort of overutilization of resources described in the tragedy of the commons.

### Simulation reveals the impact of group characteristics on task earnings

Our group decision-making task was amenable to simulation, as we gave no information to participants about other individuals in their group or feedback as to what other group members had chosen. We were therefore able to bootstrap large amounts of randomly simulated groups by repeatedly sampling ten participants and then evaluating the results of the decisions of those hypothetical groups. We first used this analysis to determine the impact of alternative offer level and amount of resource availability on overusing the resources. After simulation, we determined that both reward level and resource availability impacted resource overuse, with higher reward levels leading to higher resource overuse and more resources available leading to lower resource overuse (Fig. 4). Reward level influenced resource overuse more than the amount of resources available, likely because increased resource availability intrinsically compensated for an increase in alternative offer decisions.

**Fig. 4.**
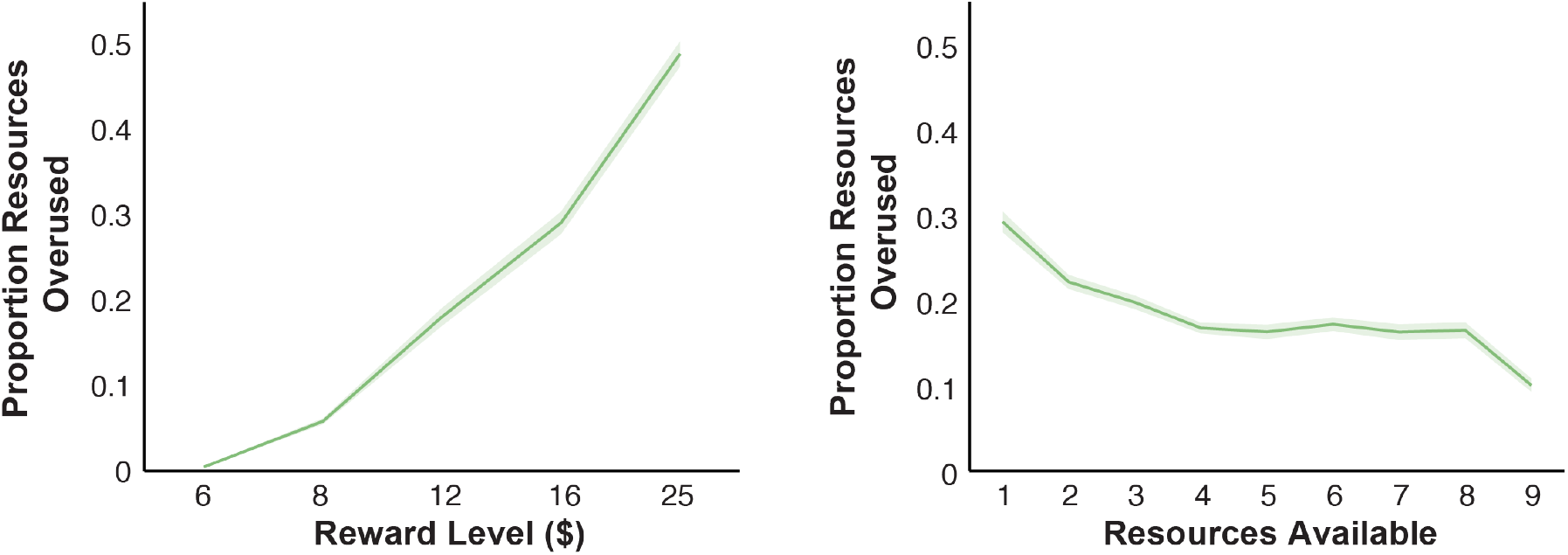
Resource overuse in the group decision-making task is impacted by both alternative offer level and amount of resources available. Proportion of resource overuse in simulated groups averaged by either alternative offer reward level (left) or amount of resources available (right). Shaded area indicates SEM.

Next, we aimed to use this simulation approach to pinpoint differences in task performance between participants in our study. For all 80 participants in our sample, we determined individual performance by repeatedly sampling nine other participants to form a group of ten with the particular individual of interest. We then evaluated the averaged individual earnings of each participant, defined as the summed total of money earned by the individual participant if all trials were to be played out, as well as the group earnings, defined as the averaged amount earned by everyone in the group other than the individual of interest. We first determined a tradeoff, demonstrating a robust negative correlation between individual and group earnings in our sample (Fig. 5; *R* = -0.957, *P* < 0.001, Spearman’s rank correlation).

**Fig. 5.**
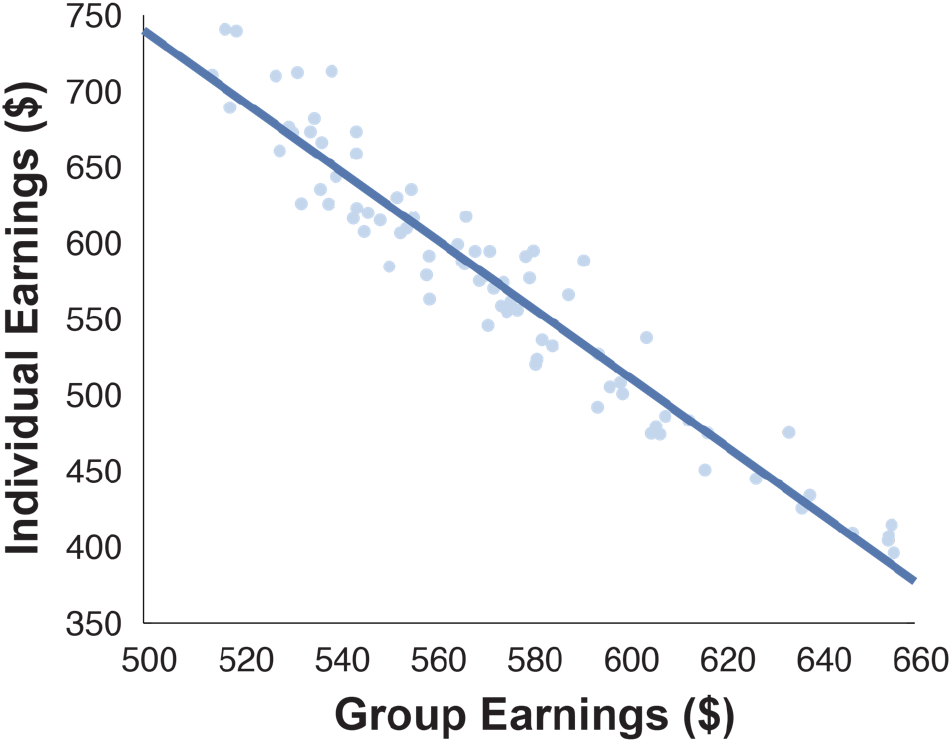
A trade-off between individual and group earnings in the group decision-making task. Individual earnings are plotted against averaged earnings of other group members for all participants. The solid trend line indicates that a significant relationship was observed (*P* < 0.001, Spearman’s rank correlation). *N* = 80 participants.

We then separated participants into two groups: “low earners,” which were defined as the bottom 30 individual earners, and “high earners,” which were defined as the top 30 individual earners. As expected, when the individual and group earnings of the two groups were evaluated in a two-way ANOVA with group and earnings type as factors, there was a main effect of low versus high earner group (Fig. 6A; *F*_1,59_ = 60.9, *P* < 0.001) as well as an interaction between low versus high earner groups and individual versus group earnings (Fig. 6A; *F*_1,59_ = 349.3, *P* < 0.001). When examining the differences in behavior in low versus high earning groups, low earners tended to choose the alternative offer less (Fig. 7; *z* = 4.78, *P* < 0.001, Wilcoxon signed-rank test) and displayed higher parabolic *k* parameters in our behavioral model (Fig. 7; *z* = 4.78, *P* < 0.001, Wilcoxon signed-rank test), indicating that low earners were more sensitive in their subjective valuation of the alternative offer when the amount of available resources were low. Together, these results indicate that high earners in our task were generally more willing to utilize resources and thus jeopardize the earnings of other group members, while low earners continued to be biased for the baseline offer that did not jeopardize earnings of other group members.

**Fig. 6.**
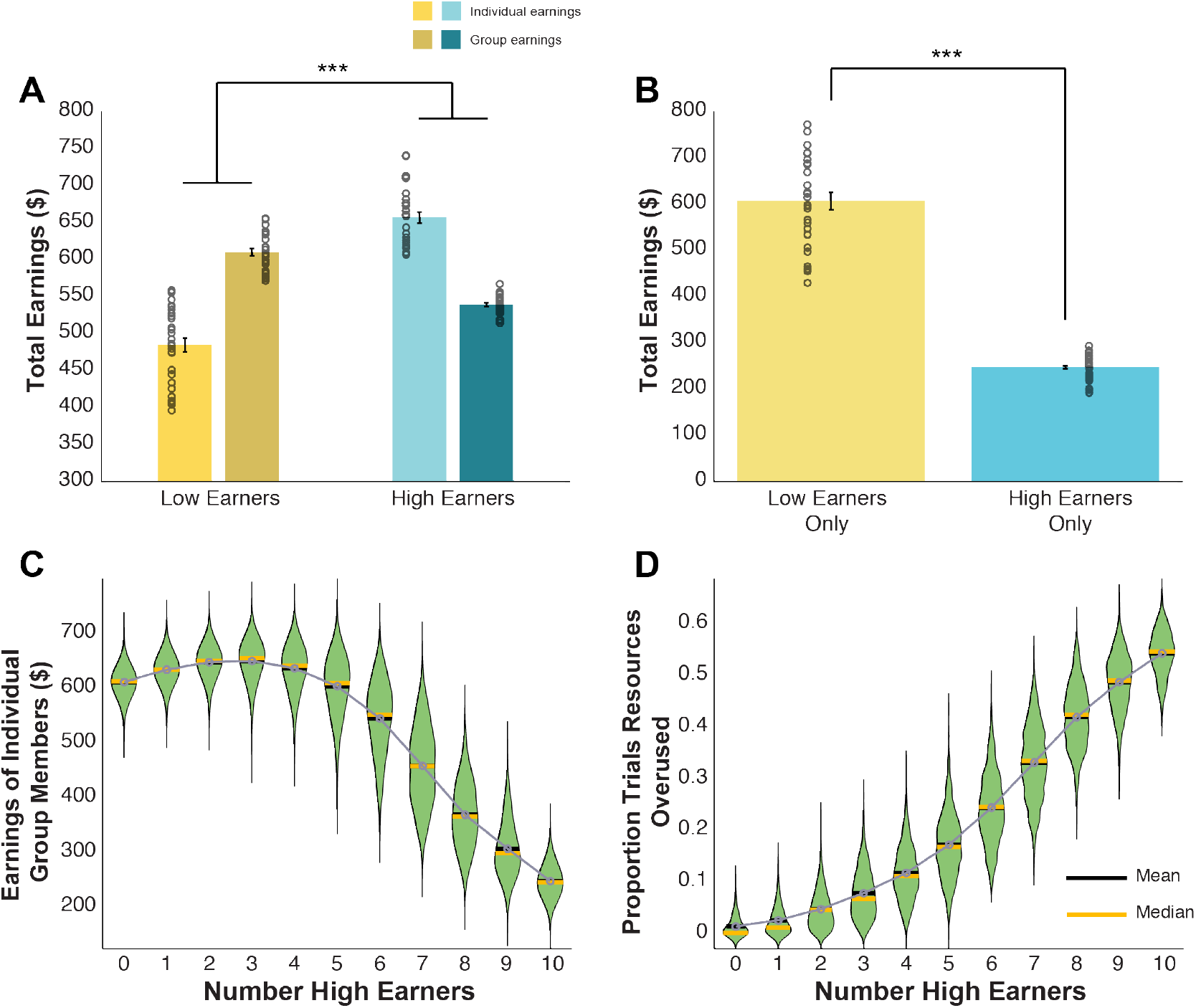
Simulations reveal divergent patterns of earnings depending on individual behavior and group composition in the group decision-making paradigm. *(A)* Total individual (light colors) and group (dark colors) earnings for low (yellow; *N* = 30 participants) and high (blue; *N* = 30 participants) earning individuals. *(B)* Total individual earnings when low earners were placed in groups consisting exclusively of other low earners (yellow) and high earners were placed in groups consisting exclusively of other high earners (blue). *** indicates *P* < 0.001, two-way ANOVA or Wilcoxon signed-rank test. *(A-B)* Error bars indicate SEM. Individual participant data points are displayed as dots. *(C)* Averaged earnings of individual group members in simulated groups as a function of progressively including more high earners. *(D)* Averaged proportion of trials in which resources were overused in simulated groups with increasing number of high earners included. *(C-D)* Means are indicated by black lines and connected in grey while medians are indicated by orange lines. Shaded green areas indicate the full distribution of simulated outcomes using a kernel density estimation with a bandwidth of 20 *(C)* or 0.01 *(D)*.

**Fig. 7.**
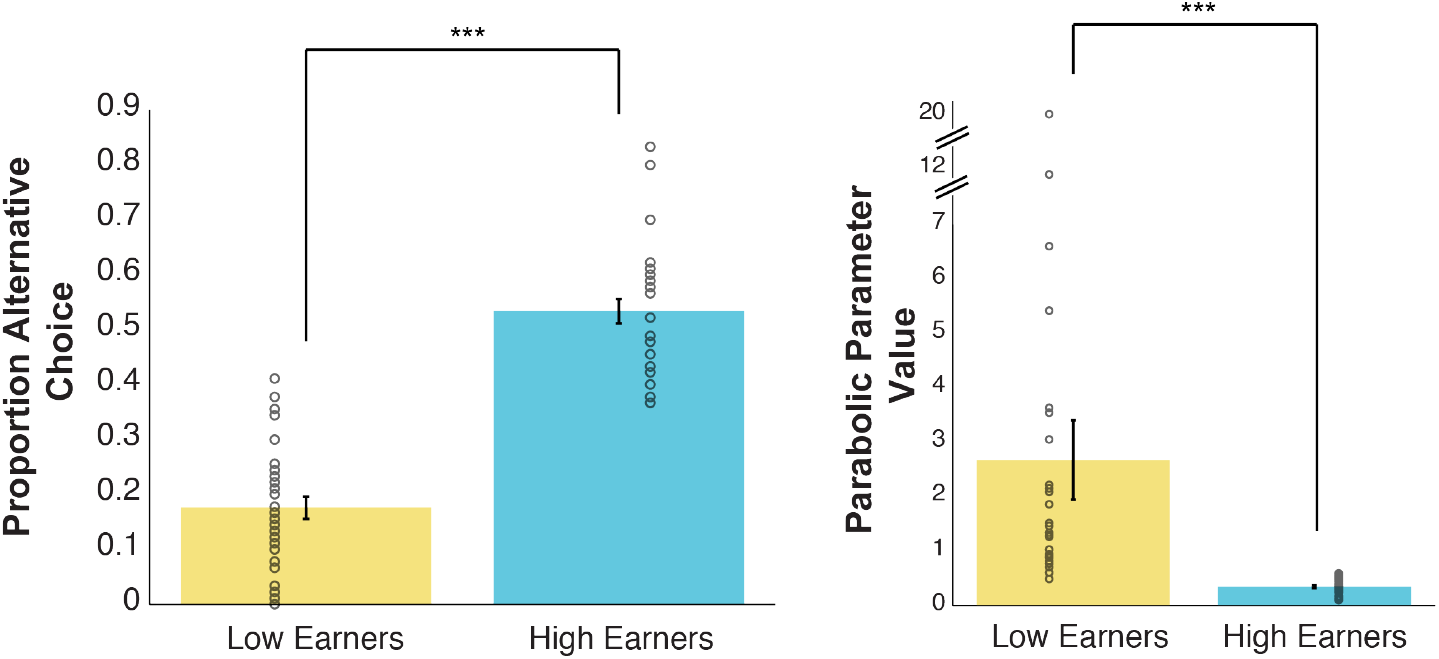
High earners are more willing to utilize resources than low earners. Proportion of alternative offer choices are displayed for low (yellow) versus high (blue) earners on the left. Parabolic parameter values (*k*) for low and high earners are displayed on the right. Error bars indicate SEM. Individual participant data points are displayed as dots. *** indicates *P* < 0.001, Wilcoxon signed-rank test. *N* = 30 low earners and 30 high earners.

While high earners were more successful individually when placed in groups that were mixed with low and high earning group members, we next aimed to determine whether high earners would still outcompete low earners when groups were homogenous in that they consisted solely of either low or high earners. For this analysis, we evaluated the individual earnings of all low and high earners, but this time we placed them in groups that consisted exclusively of other low earners or exclusively of other high earners, respectively. This yielded a dramatic reversal of individual earnings, such that low earners in groups consisting only of other low earners earned more compared to high earners in groups consisting only of other high earners (Fig. 6B; *z* = 4.78, *P* < 0.001, Wilcoxon signed-rank test). Finally, we also constructed groups of varying compositions, progressively ranging from groups with no high earners to groups consisting only of high earners. Across group compositions, we evaluated both the averaged summed earnings of all members in the group as well as the proportion of trials in which resources were overused. As more and more high earners were added to the group, group earnings dramatically decreased (Fig. 6C) and resource overuse markedly increased (Fig. 6D), suggesting that this decrease in averaged group earnings as more high earners were added to the group was likely driven by increasingly prevalent resource overuse.

### Behavior in the group decision-making task correlates with self-reported social attitudes

We next examined whether people’s social attitudes were related to how they participated in our group decision-making task using three questionnaires related to altruism, empathy, and psychopathy, which were completed by the participants at the end of the study. To consolidate these scores into a single measure of social attitudes, we performed a principal components analysis on the scores from all 80 participants in our sample (Ereira, Dolan, & Kurth-Nelson, 2018). The first principal component explained 61.3% of variance and loaded negatively onto altruism and empathy (altruism coefficient = –0.35, empathy quotient = –0.60) and positively onto psychopathy (Fig. 8A; psychopathy coefficient = 0.72), indicating that this first principle component was primarily a measure of antisocial attitudes. We observed a negative relationship between the antisocial principle component score and the parabolic *k* parameter from our behavioral model (Fig. 8B; *R* = -0.348, *P* = 0.004, Spearman’s rank correlation). This indicates that those with higher measures of antisocial attitudes were less sensitive in their subjective valuation of the alternative offer depending on the amount of resources available, in turn likely resulting in their resource overuse. Finally, there was a trend for high earners to have higher antisocial attitude component scores than low earners (Fig. 8C; *z* = 1.70, *P* = 0.090, Wilcoxon signed-rank test). These results together indicate that various components of our task corresponded broadly with antisocial personality measures.

**Fig. 8.**
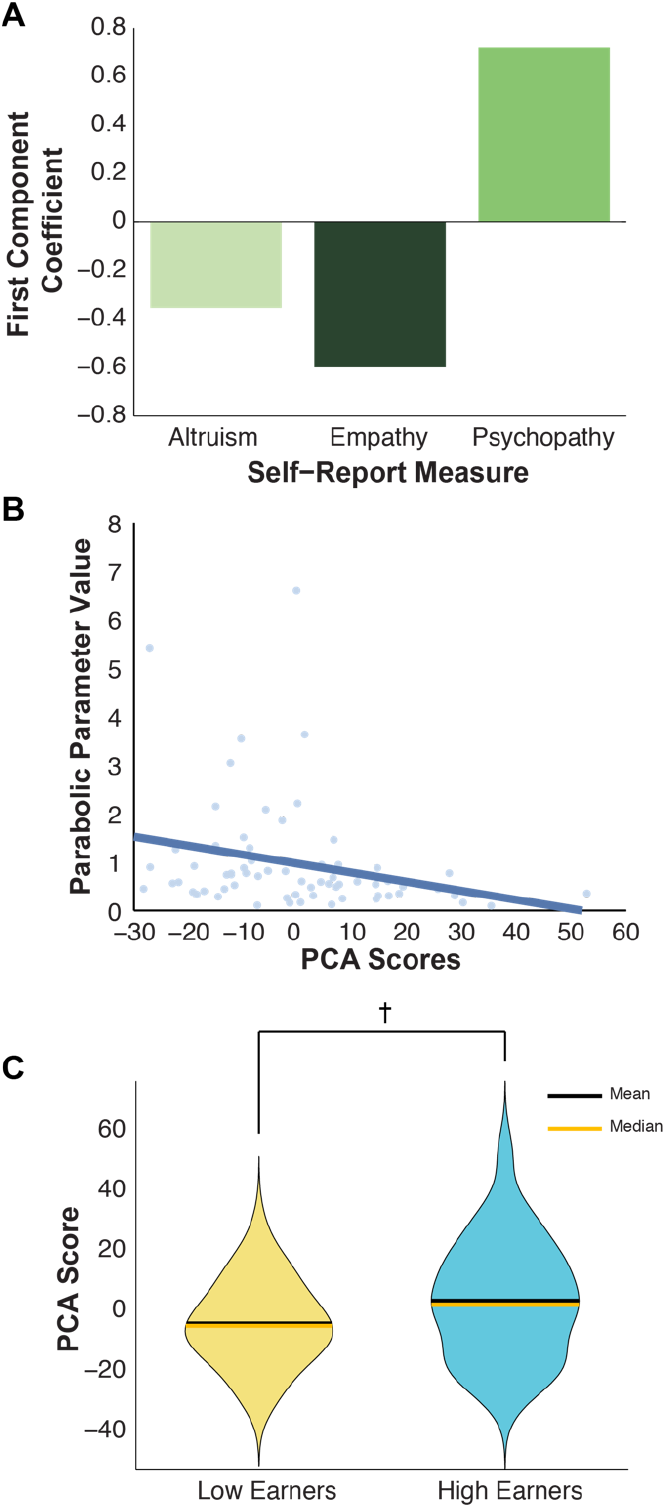
Individual differences in social attitude measures are related to behavior in the group decision-making paradigm. *(A)* Correlation coefficients for the first principal component of a principal component analysis of altruism, empathy, and psychopathy scores. *(B)* The parabolic parameter value (*k*) for each participant is plotted against the first principle component score. The solid line indicates a significant relationship (*P* < 0.05, Spearman’s rank correlation; *N* = 70 participants). *(C)* A trend was observed for high earners to have higher first principal component scores than did low earners. Means are indicated by black lines while medians are indicated by orange lines. Shaded areas indicate the full distribution of individual scores using a kernel density estimation with a bandwidth of eight. † indicates *P* < 0.1, Wilcoxon signed-rank test. *N* = 30 low earners and 30 high earners.

## DISCUSSION

In this study we used a new approach to study and informatively simulate the tragedy of the commons scenario in a computerized environment. Data collected via the personal computing devices of 80 participants yielded multiple insights into group decision-making concerning scarce, commonly held resources. As expected, participants adjusted their willingness to utilize resources depending on the potential increase in monetary gain from using resources as well as the amount of resources available. Participants were found to follow a parabolic decision-making process in determining their subjective valuation of resource usage depending on the amount of resources available, stipulating that the utilization of resources was strongly discounted only in trials in which exceptionally few resources were available. Simulation analyses furthered understanding of individual decision-making by revealing a tradeoff between individual and group earnings, such that participants varied in their willingness to use resources for their own benefit and in turn to the detriment of unfamiliar others in the assigned group. Finally, willingness to utilize resources was found to correlate with a composite measure of self-reported social attitudes. Together, these findings support the importance of the behaviors and attitudes of individual group members in influencing group outcomes.

In particular, the present results shed light on a central issue in the study of decision-making in the social sciences, namely whether or not groups invariably overuse commonly held resources (Berkes, et al., 1989; Boyd, et al., 2018; Feeny, et al., 1990). Our findings generally indicate that this is not the case. In randomly simulated combinations of participants, resource overuse only occurred on average in approximately 20–25% of trials, although resource overuse was much more prevalent in trials with higher alternative monetary values (Fig. 4). Instead, the current results strongly support the importance of group composition in the extent of resource overuse. To date, the literature regarding the tragedy of the commons has largely treated groups of similar size and similar amounts of commonly held resources as equally likely to overuse these resources. The current findings instead emphasize the importance of the decisions made by individual group members, and thus their attitudes, in determining group outcomes. Fascinatingly, in the sample of 10-person groups analyzed, a group composition of three high earners and seven low earners led to maximal averaged group earnings (Fig. 6C). Although averaged resource overuse increased slightly as zero to three high earners were included in simulated groups (Fig. 6D), the amount of resources able to be extracted by high earners, who were generally more likely to use resources, likely counteracted any added resource overuse. However, as more than three high earners were included in simulated groups, averaged group earnings catastrophically declined, supporting the presence of a critical tipping point dictated by group composition (Fig. 6C). This effect was likely due to differences in resource overuse between groups of varying individual compositions. For instance, median resource overuse was 0% in groups consisting only of low earners, whereas groups consisting only of high earners overused resources in more than 50% of trials (Fig. 6D).

Computerized paradigms amenable to emulating the tragedy of commons such as the one developed and used in this study will be fruitful in a multitude of further applications. While we observed interesting individual differences relating to social attitudes (Fig. 8), other attitudes should also play an important role in underlying behavior in the group decision-making task. Specifically, further analyses concerning the effect of other individual differences such as risk and ambiguity aversion with respect to resource depletion as well as impulsivity may critically elucidate the fundamental psychological underpinnings of decisions in our paradigm. Furthermore, other individual factors, such as income and political affiliation, as well as social reputation sensitivity (Izuma, Matsumoto, Camerer, & Adolphs, 2011; Meshi, Morawetz, & Heekeren, 2013), an ability to predict others’ minds (Tamir & Thornton, 2018), and in-group/out-group compositions (Banaji, Baron, Dunham, & Olson, 2008), would be particularly illuminating in the context of group decision-making. The convenient dissemination of the group decision-making task presented here also allows for the rapid testing of various task manipulations designed to decrease resource overuse. Such manipulations could include direct or indirect feedback concerning the decisions of other group members or a system of imposed second- or third-party punishment for those who contribute to resource overuse.

Research inspired by economics and psychology in the context of neuroimaging have yielded fundamental insights into the human brain, including determining the neural correlates of action selection (Daw, O’Doherty, Dayan, Seymour, & Dolan, 2006), intertemporal choice (Kable & Glimcher, 2007; Lin, Saunders, Hutcherson, & Inzlicht, 2018), risk and ambiguity aversion (Hsu, Bhatt, Adolphs, Tranel, & Camerer, 2005; Huettel, Stowe, Gordon, Warner, & Platt, 2006), decision-making involving others (Hutcherson, Bushong, & Rangel, 2015; Nicolle et al., 2012; Sul et al., 2015), and certain mental disorders (Kishida, King-Casas, & Montague, 2010). Determining how information regarding individual and group reward signals are integrated in the human brain in the context of scarce resource allocation will be noteworthy and may also help to broadly inform our understandings of human cooperation (Rand & Nowak, 2013) and altruism (Fehr & Fischbacher, 2003). Further work using a similar paradigm like the one presented here may continue to provide important mechanistic insights into the tragedy of commons that have implications for everything from everyday decision-making to national and international policies regarding shared resources. For now, the present results strongly indicate the importance of individual group composition on shared outcomes.

## ACKNOWLEDGMENTS

We thank Spencer Birney for his help in testing a pilot version of our experimental task. We also thank Amrita Nair for her administrative assistance throughout this study. Funded by a Natural Sciences and Engineering Research Council of Canada Fellowship (PGSD3-471313-2015) awarded to M.P.

## AUTHOR CONTRIBUTIONS

M.P., I.L., and S.W.C.C. designed experiments. M.P., S.A., and R.J. piloted the study and collected all data. M.P. and S.A. analyzed all data and developed all behavioral models. M.C., M.P., S.A., R.J., I.L., and S.W.C.C. wrote and edited the paper.

